# Using the *E. coli* Alleleome in Strain Design

**DOI:** 10.1101/2023.09.17.558058

**Authors:** Patrick Phaneuf, Zofia D. Jarczynska, Vijayalakshmi Kandasamy, Siddharth Chauhan, AM Feist, Bernhard O. Palsson

## Abstract

Leveraging observed variants in strain design is a promising technique for creating strains with specific properties. Adaptive laboratory evolution (ALE) experiments generate variants that enhance fitness under specific conditions and can contribute to application-specific strain designs. Further, the wild-type (WT) coding alleleome of an organism, the complete set of its genes’ WT alleles, can provide an additional amount and diversity of variants not yet accessible from the aggregation of ALE experiment results. This study used both an ALE mutation database (3093 genomes) and a large set of WT genomes (12,661 genomes) to explore the sequence solution space of genes involved in tolerance to 10 conditions of industrial importance. To accomplish this, ALE variants for 22 genes previously identified as potentially important for industrial chemical tolerance were collected and supplemented with all available variants from the WT coding alleleome. A total of 4879 variants were reintroduced and used in 10 selection experiments. Both ALE and WT contributed highly enriched variants, where the enrichment and benefits depended on the conditions, genes, and gene product regions. The results also revealed that variants not originating from the initial experiment could potentially confer substantially greater benefits. Additionally, ALE and WT variants rarely overlapped on AA positions, but their clustering did coincide with where highly enriched variants were ultimately located. For genes primarily hosting potential gain-of-function variations, substitutions predicted to have a conservative impact frequently outperformed more radical substitutions. Case studies demonstrated that maximizing the amount of variants enabled easier identification of variant trends, which in turn can be used to better understand areas and characteristics of genes that can be feasibly varied, representing what could be thought of as a genome design variable. The combination of ALE and WT variants is a promising approach for use in future projects to better constrain and ultimately achieve practical coverage in the exploration of feasible sequence solution space.

**Visual Abstract:** 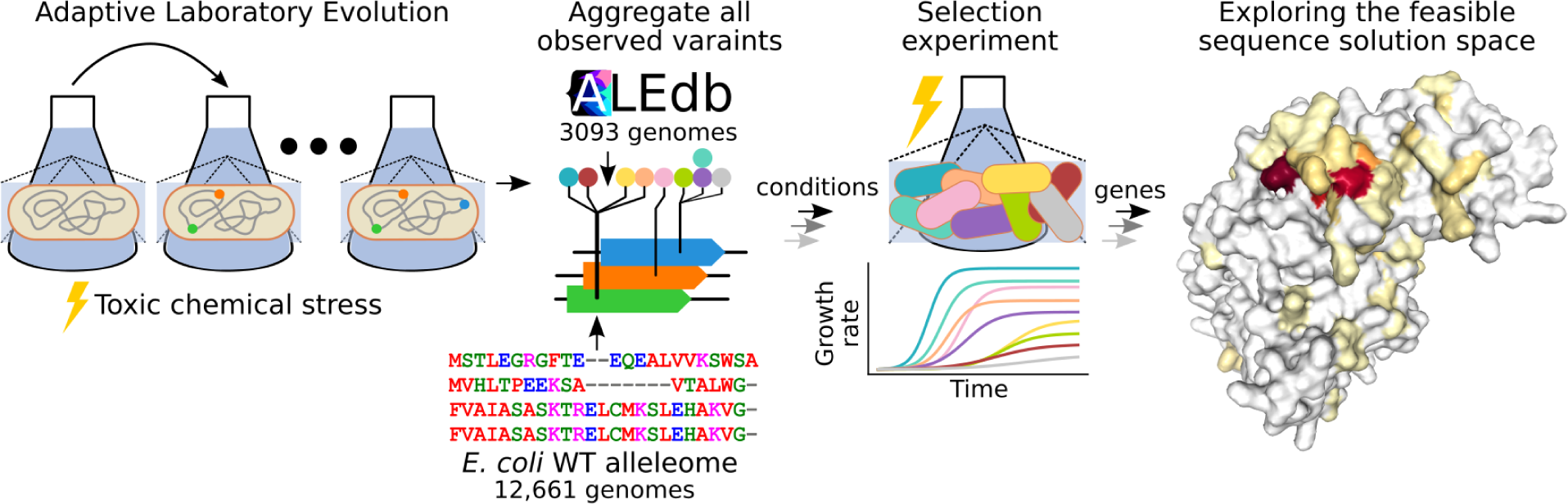

## Introduction

In the burgeoning field of microbial genomics, Adaptive Laboratory Evolution (ALE) has emerged as a critical tool for illuminating the genetic underpinnings of environmental adaptation. By applying selective pressure over multiple generations, ALE experiments generate diverse genetic variants that enhance organismal fitness under specific conditions, a feature of particular relevance to applications in biotechnology and medicine (1,2).

Recent research has demonstrated the value of maximizing variants to genomic features of interest to understand functional changes that beneficial variants converge on (3,4). The field of pangenomics research has been aggregating genomes of numerous species isolated from a variety of environmental and clinical conditions, generally referred to as wild-type strains (WT), and developing methods to compare their gene sets and alleles (5,6). WT strains contain sequence variants selected for by pressures from their natural environments and these variants can be used in strain design, much like the mutations derived from ALE experiments. *Catoui et al.* had also shown that WT strains can provide many more variants than ALE strains (WT: 503,744 variants / 2,661 strains = 189 AA variants per strain; ALE: 25,470 variants / 4,181 strains = 6 AA variants per strain) (6). The variants found across WT alleles can thus provide a scale of variants not yet accessible solely from the aggregation of ALE experiment results (6). Additionally, ALE and WT variants have demonstrated little overlap, complementing each other, and therefore may cover the potential feasible functional solution space of a gene better than variants acquired from ALE alone (6).

Previously, the reintroduction of specified variants at a large scale was inaccessible. Recent technologies that precisely deliver single edits into the genome of individual strains have enabled the parallel introduction of up to ten thousand mutations per pool, enabling the exploration of their phenotypes (7). This study leveraged these methods to compare the fitness benefits and characteristics of variants from a comprehensive *E. coli* coding alleleome comprising of variants from both ALE and WT coding alleles. Key mutated genes from an industrial chemical tolerance (ICT) ALE study (8) were chosen as variant targets to explore the value of natural variants for strain designs involving industrial biotechnology. Selection experiments using the selection pressures from the original ALE study (i.e., chemical tolerance) were used to derive phenotypes for each variant. By exploring the phenotypes and property changes for the potential gain-of-function variants selected from both ALE and WT sources, the functional solution space for these key genes could be characterized, thus deriving potential genome design variables.

## Results

### Dataset size and dimensions

To identify genomic features and their variants for reintroduction, the results from the ICT ALE experiments were analyzed to find potentially beneficial mutations. The original ICT ALE experiments were adapted for tolerance to toxic concentrations of an industrially relevant chemical from a set of 11 different chemicals (8), where 10 of the experiments resulted in relatively clear outcomes. Genomic features hosting potentially beneficial mutations were identified by observing the amount of replicate ALEs a genomic feature was mutated in; if a genomic feature is mutated in a large proportion of replicate ALEs, it is said to demonstrate convergence (4) (Figure 1A).

**Figure 1.**
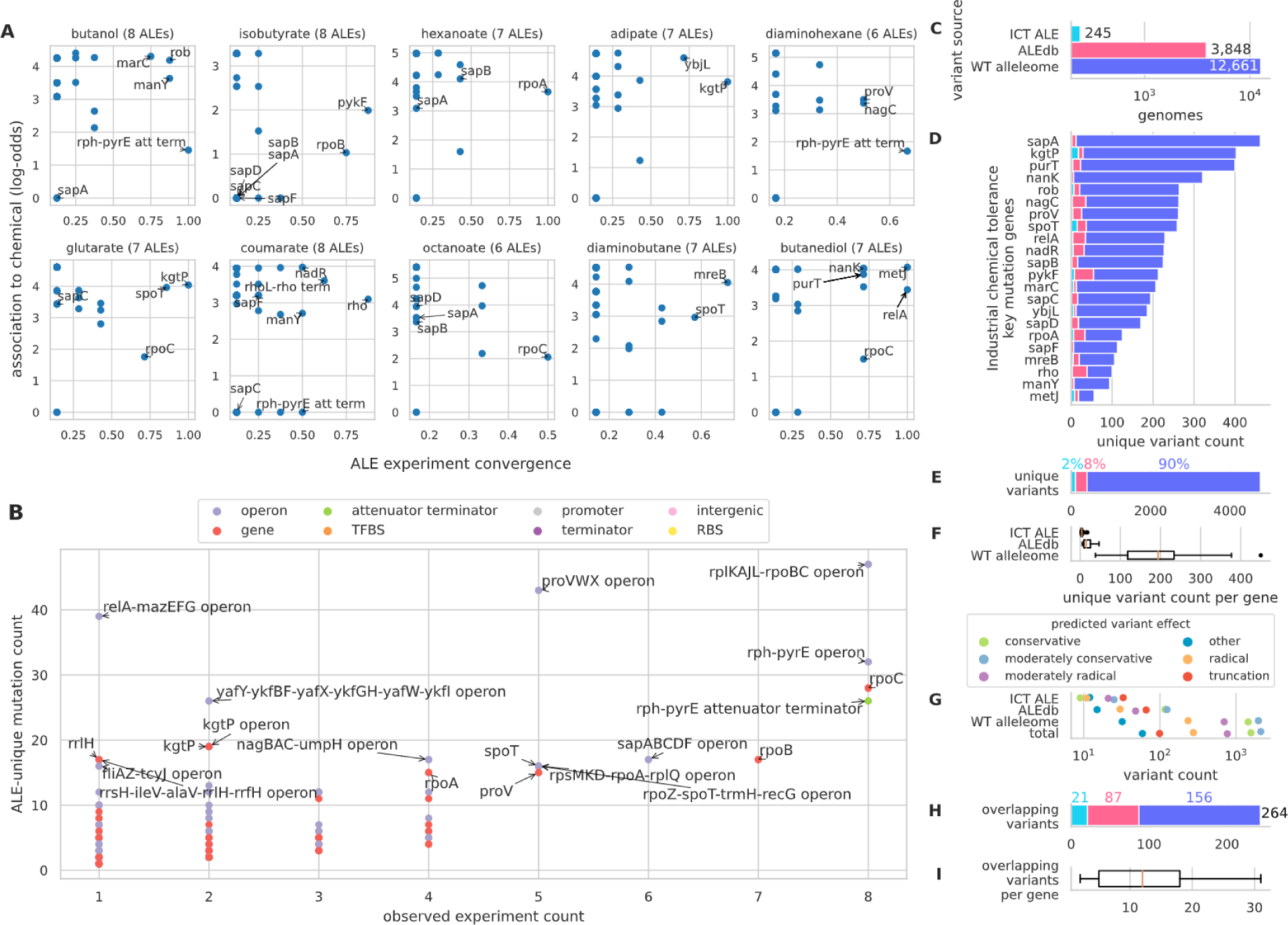
Variant targets for reintroduction. **A)** Mutated genomic features from the original ICT ALE study, where those features with convergence >= 0.5 and significantly associated are annotated along with mutated genes belonging to the sapABCDF operon. This analysis defined the target set of features to explore with additional variants. See Methods for convergence function. **B)** Count of experiments each feature was found to be mutated in. **C)** The amount of samples for each variant source. The colors of this plot are re-used throughout the manuscript to represent these variant sources. **D)** The amount of unique variants extracted from multiple sources for 22 genes of interest. **E)** The total amount of variants available for reintroduction. **F)** The distribution of variants available for each gene based on their source. **G)** The amount of variants per source per predicted variant effect. See Methods section for predictive variant effect methods. **H)** Non-truncating amino acid substitutions from either an ALE or WT source that share overlapping positions with the other source. **I)** The distributions for the amount of overlapping variants per gene.

In addition to convergence, mutated features were also statistically associated with conditions to understand which mutated features were potentially beneficial for conditions of interest (4) (Figure 1A). Some individual genomic features were mutated at low frequencies across ICT ALE experiments, though their operons were frequently mutated, indicating that the systems in which they encode were important targets for mutations. The SapABCDF operon, encoding the SapBCDF complex implicated in putrescine export (9), is one such example (Figure 1B). The combination of convergence, statistical associations to conditions, and frequency of mutation across multiple ALE experiments involving tolerance to toxic chemical concentrations highlighted a subset of genomic features potentially hosting beneficial variants.

According to these methods, 22 genes were selected from across the original ICT ALE experiments for the reintroduction of variants. The variants were aggregated from two sources: a database of ALE experiment mutations (10) and an E. coli wild-type coding alleleome (WT alleleome). The WT alleleome was derived from a database of wild-type E. coli genomes (11) (Figure 1C, 1D). In total, 4879 unique variants were aggregated with 90% coming from the WT alleleome (Figure 1E) which generally contributed 10 times more variants to genes than ALE sources (Figure 1F).

To develop better intuition as to the potential impact of these variants, analysis was performed to predict their effects on the translation of a gene. These predictions are based on Grantham score categories for AA substitutions (conservative ≤ 50, 51 ≤ moderately conservative ≤ 100, 101 ≤ moderately radical ≤ 150, radical ≥ 151) (12) and if variants frameshifted or truncated the open reading frame of a gene (grouped together as a truncation). The results showed that across all variant sources, moderately conservative and moderately radical substitutions consistently have a high abundance, with radical having mid-to-low frequency (Figure 1G). Variants in the “other” category, which is mostly comprised of in-frame insertions and deletions, were found in low amounts for all sources. Conservative substitutions were frequent with larger variant sources and truncations frequent with ALE variant sources.

The overlap of variants from the different sources of variation was investigated. Overlapping variants had been observed as rare in a broad study of ALE vs WT variants, suggesting that they may indicate possible important outcomes for these variants (6). A total of 264 non-truncating amino acid substitutions were found to overlap on positions with a variant from another source (ALE vs WT) (Figure 1H). Additionally, all genes hosted at least one of these overlapping variants (Figure 1I).

### Enrichment of reintroduced variants

Experimental validation was performed to compare the impact of WT and ALE variants for the target genes found via ALE experimentation. All variants were successfully reintroduced into a population (see the Sequencing of Library Samples subsection of the Methods section) and samples from this population were subjected to selection experiments of stressful concentrations with the 10 targeted industrial chemicals. Cultures grew within 8 of the 10 conditions and the variant abundances were measured using barcode amplicon sequencing (See Methods).

Broadly, all three sources provided statistically enriched variants for the treatments (see the Enrichment Scores subsection of the Methods section) (Figure 2A) with rankings that generally coincided with the overall amount of variants each source contributed (Figure 1E): WT provided the most amount of input variants and highly enriched variants, followed by ALEdb and ICT ALE. Either ICT ALE or ALEdb provided variants with generally higher enrichment (median) and maximum enrichment in 5/8 conditions, and WT provided generally higher enrichment (median) in the remainder (Figure 2A, Figure S1A, Figure S1B). When considering the individual genes per condition, every variant source provided a variant with the highest enrichment (Figure 2C). In all conditions, there were genes with highly enriched variants only provided by WT or ALE, though more often WT (Figure 2C). In specific conditions, key genes *spoT* (diaminobutane, glutarate), *sapA* (octanoate), and *sapB* (octanoate) only hosted highly enriched variants from WT. The variant of highest enrichment for *metJ* was from WT in diaminobutane but from ALE in butaindiol. For *rpoA*, ALE provided the highest enriched variant for butanediol, diaminobutane, glutarate, and hexanoate, WT provided the highest enriched variant for octanoate (Figure 2C), and both ALE and WT provided closely tied variants for the highest enrichment on diaminohexane (Figure 2C). These results indicate that the WT variant source offers not just valuable variants, but those with the highest benefit.

**Figure 2.**
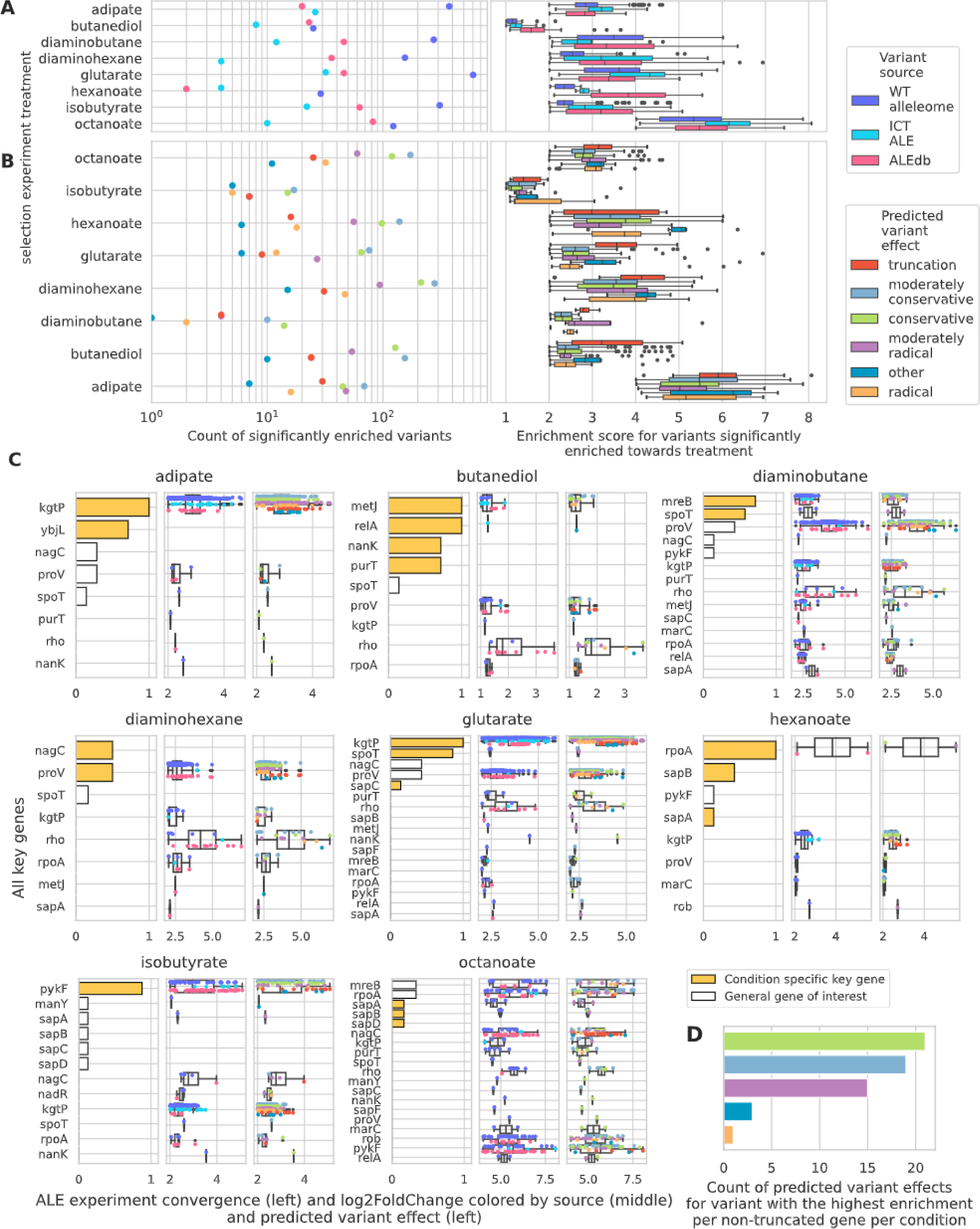
A comparison between enrichment of ALE versus WT variants on multiple levels. Condition-dependent score filtering was used to better focus on genes with high-scoring variants. **A)** The amount and enrichment score distributions for highly enriched variants according to chemical selection pressures and variant sources or **B)** predicted variant effects. **C)** Enrichment score distributions for highly enriched variants across conditions. ALE experiment convergence values indicate how many individual ALEs a gene were mutated in an ALE experiment. Genes with 0 convergence values designate genes that weren’t considered key within their original ALE experiments. **D)** The count of predicted variants effects for variants with the highest enrichment per non-truncated gene per condition.

The ranking of variants by count according to predicted mutational effects was largely consistent when comparing variants significantly enriched per condition (Figure 2B) and variants present prior to selection experiments (Figure 1G). Variants expected to truncate or have “other” (mostly in-frame INDELs) effects on genes generally granted slightly higher enrichment, with all AA substitutions having similar general phenotypic distributions (Figure S1C), where conservative, moderately conservative, or moderately radical (and never radical) were seen to achieve the highest enrichment in 75% of the conditions (Figure 2B, Figure S1C). For non-truncated genes, the variants that provided the most enrichment were characterized as conservative, moderately conservative, and moderately radical AA substitutions (Figure 2D). While WT amino acid substitutions at sites that overlap with ALE variant sites were hypothesized to have a greater phenotypic effect in ALE conditions compared to non-overlapping substitutions (6), this study found no evidence to support this hypothesis (Figure S1B).

Those genes with medium to high levels of convergence from ALE experiments also often demonstrated variants enriched more highly for the treatment than most other genes, supporting the expectation that ALE-selected mutant genes (condition-specific key genes) should be highly enriched (Figure 2C, Figure S1D). This wasn’t the case for all genes mutated in ALE experiments, potentially due to their ALE mutations being neutral or because they required other mutations to exist before becoming beneficial (Figure 2C, Figure S1D). Also, some genes not mutated in specific ALE experiments hosted highly enriched variants in selection experiments (Figure 2C, Figure S1D); for example, *rho* variants were highly enriched in most selection experiments though was only a key mutated gene in ALE experiments involving coumaric acid. These deviations from ALE experiments could indicate epistatic relationships with other mutants in ALE strains that weren’t reintroduced in this study’s single variant strains.

Highly enriched variants were expected to more clearly describe functional information about targets than variants of low enrichment and were therefore the focus of further analysis. Highly enriched variants were chosen according to the variant enrichment distributions per condition (Figure 2).

### Case studies

A set of genes were selected for a deeper comparison of variant properties between ALE and WT variants. The genes selected were according to if they were deemed as key genes from ALE experiments and hosted highly-enriched and mostly non-truncating variants. Those genes were *rho, rpoA,* and *metJ*.

### Rho

*rho* variants were highly enriched in ⅞ of the selection experiments (Figure 2C). Additionally, none of the specific variants enriched were truncations, indicating benefit from possible gain-of-function effects rather than the disruption of the gene product’s overall functionality. Rho had additionally been identified as a key mutated gene with the original ICT ALE experiment involving high concentrations of coumarate, though final concentrations didn’t yield any substantial growth with this work’s selection experiments. Rho provides one of two forms of transcription termination with *E. coli* by forming a hexamer that binds with nascent RNA transcripts and halting transcription through template DNA displacement (13).

Variant clustering was investigated on the amino acid sequence and the 3D structure (Figure 3). Variants were generally spread across the sequence though there were hotspots (Figure 3A). Variants between WT and ALE sources infrequently overlap on positions (Figure 3A), though the hotspots did host specific positions with substantial overlap. These hotspots also coincided with the highly enriched variants (Figure 3A). Highly enriched variants clustered on or near the RNA binding domain (Figures 3A, 3B). These variants were selected in multiple different conditions and come from both variant sources, though sources for highly enriched variants rarely overlapped on specific residues (Figure 3B).

**Figure 3.**
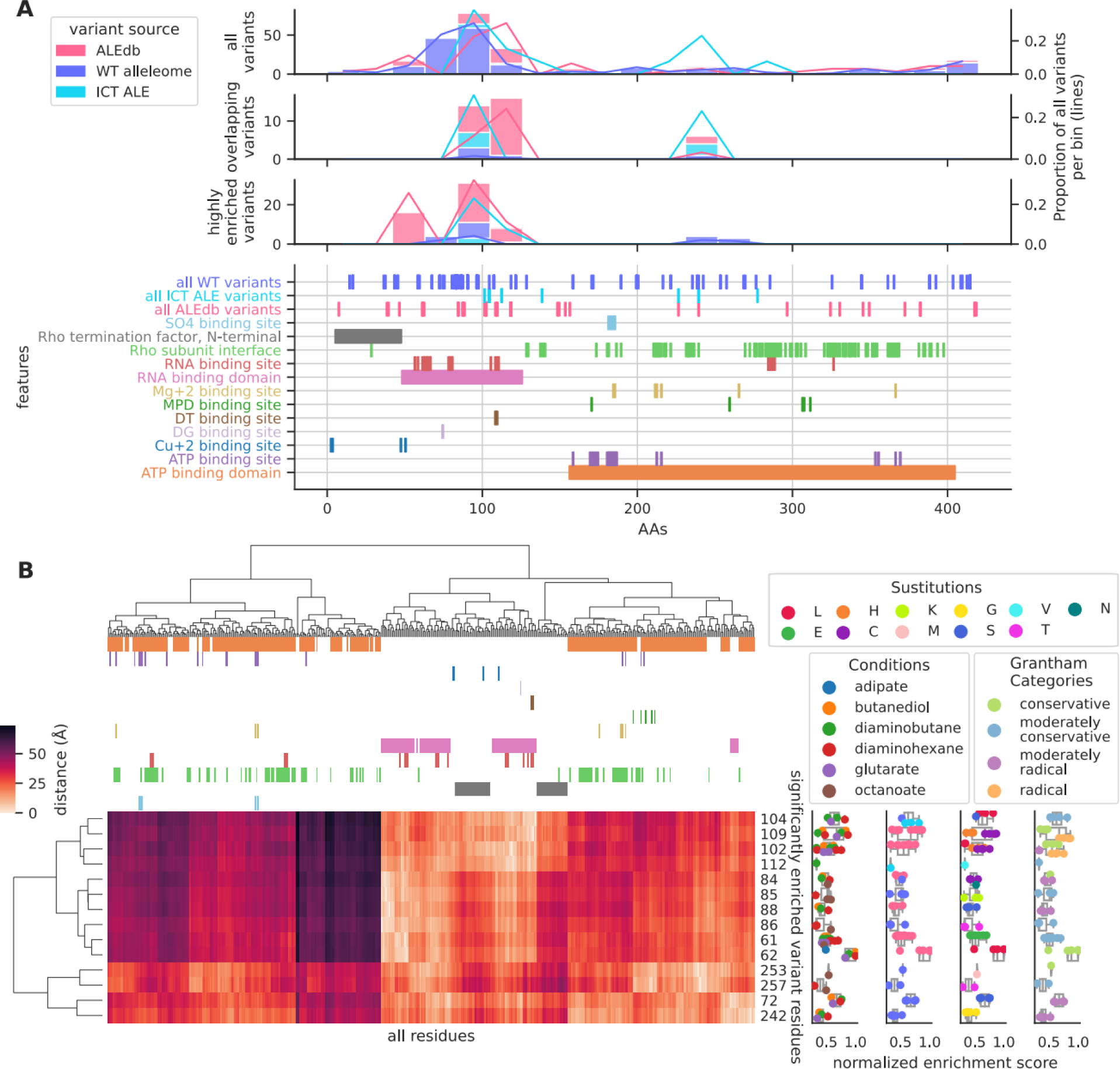
**A)** A summary of all, overlapping, and highly enriched Rho variants across the amino acid sequence and its functional annotations. An overlap is identified when a variant from either ALE source overlaps with a WT variant. **B)** Clustering of highly enriched Rho variants versus all residues according to their 3D distance on the protein structure as well as their enrichment scores and variant properties. The colors for the clustermap columns represent the same features in A) and use the same color scheme. The boxplots with colored points describe the normalized enrichment score across all conditions of each highly enriched variant for the gene and the coloring of the points describes different categories of information (1st: Conditions, 2nd: Variant Source, 3rd: Substitutions, and 4th: Grantham Categories). The coloring for the Variant Sources reuses the legend from A). Enrichment scores were normalized across conditions to account for experimental variability (see Methods).

The large variety of conditions that selected highly enriched variants on most variant residues suggested that these variants confer broad benefits for tolerance to the chemical categories used in this study. This allowed the variants to be compared with each other. ICT ALE variants provided the highest enrichment on their target residues though other ALE mutated residues reached higher enrichment (Figure 3B). WT variants provided high enrichment on the residues they targeted, though generally had lower enrichment than ALE mutations on residues targetted by both ALE and WT variants (Figure 3B, residues 84 and 104). Highly enriched ALE variants preferred variant clusters involving residues between 102 and 112 or 62 and 88 while selected WT variant clusters involved residues between 253 and 257 or 72 and 242 (Figure 3A and 3B), demonstrating that WT variants can manifest more beneficial variants than ALE in specific regions. Most variant residues had very specific substitutions, though some hosted multiple substitutions (Figure 3B, residues 84, 102, 104, 109); substitutions to these residues with more impactful Grantham Scores categories were generally more highly enriched (Figure 3B).

ALE variants preferred a subset of property changes whereas WT variant property changes were generally more varied (Figure S1). Some variants near each other (residues 102 and 109) had very similar property changes, even with the selection of multiple different substitutions, though proximity didn’t guarantee similar substitution properties (Figure S1). Side-chain size and flexibility changes were consistently chosen by both ALE and WT variants, demonstrating that the same potentially beneficial changes were available for selection with both variant sources (Figure S1). Variants to the RNA binding domain (residues 62 and 64) have previously been published and resulted in faulty RNA binding, potentially leading to the repression of biological functions whose genes rely on Rho termination due to the stalling of RNA release from RNAP (14).

### RpoA

*rpoA* variants were highly enriched in ⅞ of the selection experiments (Figure 2C). Additionally, most of the specific variants enriched weren’t truncations, indicating possible gain-of-function effects rather than the disruption of the gene product’s overall functionality. *rpoA* encodes for the alpha subunit of RNA polymerase which has two primary domains, the N and C-terminal domains. The N-terminal domain is required to assemble RNA polymerase (15) while the C-terminal domain is involved in transcription termination involving Rho and NusA (16,17).

Variant clustering was investigated on the amino acid sequence and the 3D structure (Figure 4). Variants were spread across the sequence though there were some hotspots (Figure 4A). Variants between WT and ALE sources infrequently overlap (Figure 4A), though some of the hotspots did host specific positions with overlap. These hotspots also coincided with the highly enriched variants (Figure 4A). The C-terminal domain hosted more highly enriched ALE variants compared to the N-terminal domain (Figure 4A, 4B). On the N-terminal domain, the highly enriched ALE variants formed clusters at homodimer interfaces (Figure 4B). Highly enriched WT variants were found on both termini. On the N-terminal, they tended to cluster less well around homodimer interfaces than ALE variants (Figure 4B). On the C-terminal, they were the only variant source to closely cluster around the Na+1 binding sites (Figure 4B).

**Figure 4.**
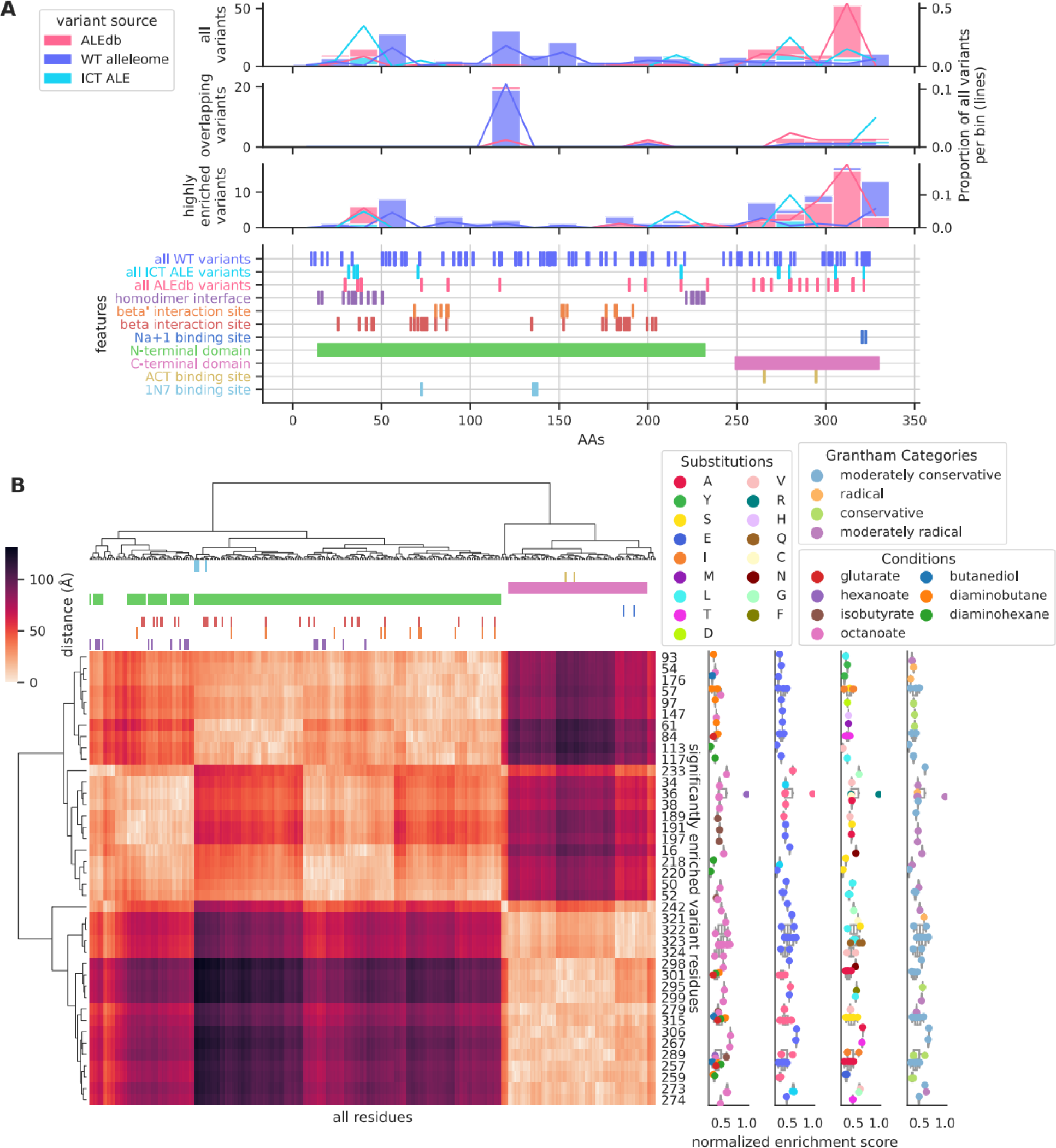
**A)** A summary of all, overlapping, and highly enriched RpoA variants across the amino acid sequence and its functional annotations. An overlap is identified when a variant from either ALE source overlaps with a WT variant. **B)** Clustering of highly enriched RpoA variants versus all residues according to their 3D distance on the protein structure as well as their enrichment scores and variant properties. The colors for the clustermap columns represent the same features in A) and use the same color scheme. The boxplots with colored points describe the normalized enrichment score across all conditions of each highly enriched variant for the gene and the coloring of the points describes different categories of information (1st: Conditions, 2nd: Variant Source, 3rd: Substitutions, and 4th: Grantham Categories). The coloring for the Variant Sources reuses the legend from A). Enrichment scores were normalized across conditions to account for experimental variability (see Methods).

Substitutions to residues were relatively specific in their substituted AA, condition, and variant source, though a subset of residues hosted up to 4 different AA substitutions from either ALE or WT variant sources (Figure 4B), providing evidence that certain residues may have multiple solutions for adjusting local function. A different set of residues hosted substitutions selected by up to 4 conditions (301, 315, and 257), though the sources and substituted AAs were unique for each residue. Excluding residues 301, 315, and 257, some conditions were relatively selective of the variant clusters (Figure 4B). Diaminobutane often selected WT variants on the N-terminal cluster that avoided homodimer interfaces (Figure 4B). Isobutyrate often selected WT variants on the N-terminal that included the homodimer interface. Octanoate mostly selected variants across the C-terminal. Substitution properties involving side-chain flexibility were also the most often selected by both variant sources, demonstrating that the same potentially beneficial changes were available for selection with both variant sources (Figure S2).

Overall, RpoA represents a significantly enriched variant target that may provide moderate benefit for most selection pressures of this study, where there is evidence of both condition-specific and general variant targets of benefit. The variants to the C-terminal may confer similar benefits as *rho* variants, and those to the N-terminal may confer benefits more specific to conditions.

### MetJ

*metJ* mutants were identified as a key mutated gene for the butanediol ALE experiment and demonstrated substantial enrichment with multiple selection experiments (Figure 2C). Additionally, none of the specific variants enriched were truncations, indicating possible gain-of-function effects rather than the disruption of the gene product’s overall functionality.

MetJ represses transcription by forming a homodimer that binds with DNA and another homodimer (co-repressor) (18). Variant clustering was investigated on the amino acid sequence and the 3D structure (Figure 5). All highly enriched variants clustered near or on repression-related functions and dimer interfaces (Figures 5A and 5B). Generally, the most highly enriched variants were often selected by multiple conditions, though sources for highly enriched variants never overlapped on residues and substitutions to each residue were unique (except with residues 6 and 39) (Figure 5B). ALE and WT variants sources both provided variants achieving the highest enrichment values for MetJ (residues 13, 20, and 27). Highly enriched variants tended to cluster near other variants of the same source, demonstrating each variant source could provide highly enriched variants for specific regions (ex: 25, 39, and 42 for WT; 13 and 14 with ALE).

**Figure 5.**
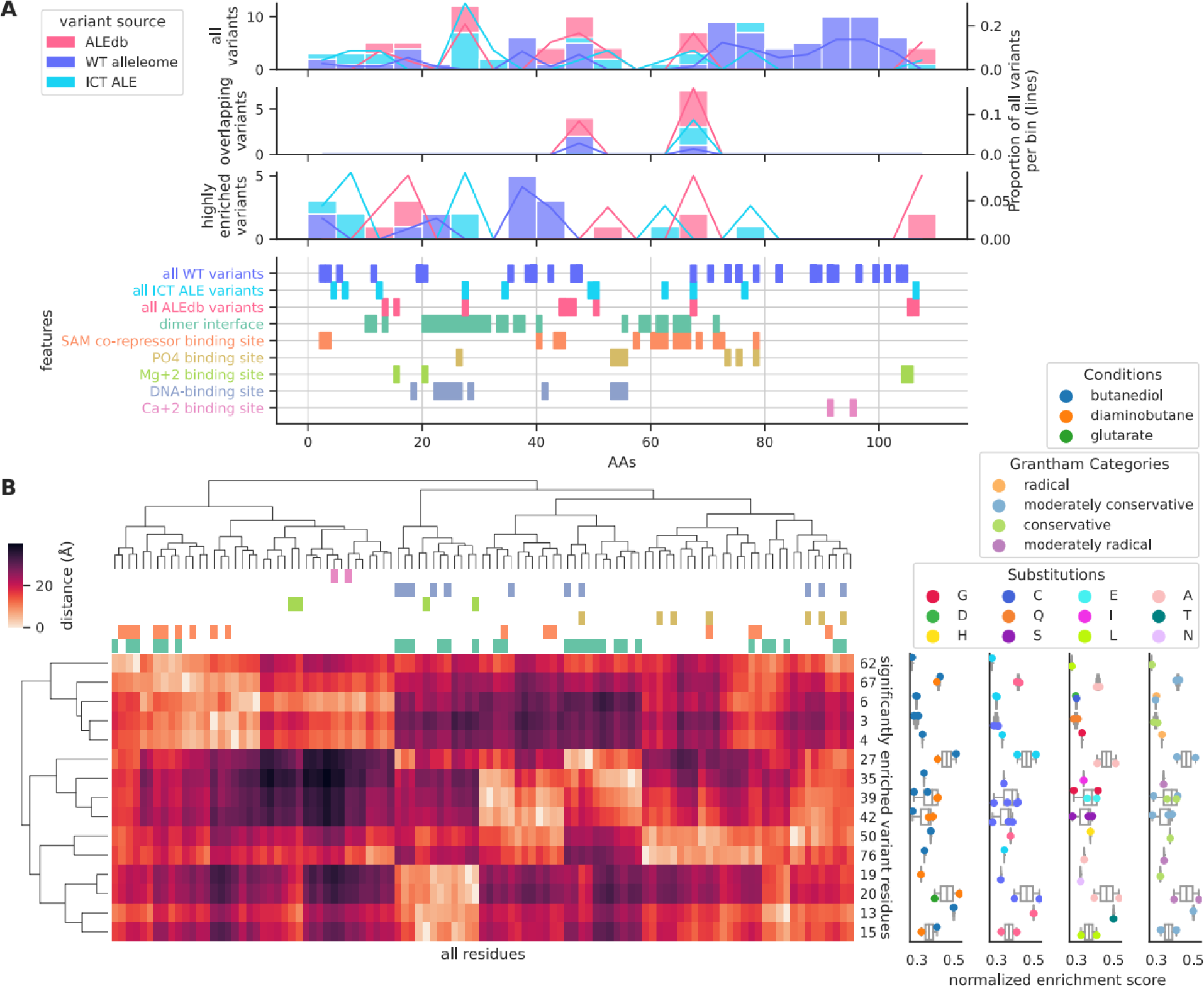
**A)** A summary of all, overlapping, and highly enriched MetJ variants across the amino acid sequence and its functional annotations. An overlap is identified when a variant from either ALE source overlaps with a WT variant. **B)** Clustering of highly enriched MetJ variants versus all residues according to their 3D distance on the protein structure as well as their enrichment scores and variant properties. The colors for the clustermap columns represent the same features in A) and use the same color scheme. The boxplots with colored points describe the normalized enrichment score across all conditions of each highly enriched variant for the gene and the coloring of the points describes different categories of information (1st: Conditions, 2nd: Variant Source, 3rd: Substitutions, and 4th: Grantham Categories). The coloring for the Variant Sources reuses the legend from A). Enrichment scores were normalized across conditions to account for experimental variability (see Methods).

Some variants in close proximity to each other demonstrated similar residue property changes (residues 13 and 15 for ALE, 35 and 39 for WT), though proximity does not guarantee similar property changes (Figure S3). ALE variants seemed to prefer the selection of side-chain flexibility changes while WT variants generally selected a larger subset of property changes overlapping with ALE (Figure S3). Variants to the co-repressor binding site and dimer interface (residue 61) have been previously published and resulted in the dimerization hindrance (18), potentially leading to increased methionine synthesis.

## Discussion

The use of variants in strain design is a promising approach to creating strains with desired properties, though finding beneficial variants for specific conditions among the vast amount of available sequence permutations is akin to finding a needle in a haystack. Experimental evolution methods have been used to reveal beneficial variants, and the aggregation of their results has made available a pool of likely-beneficial variants to leverage in strain design (4). More recently, the field of pangenomics has demonstrated access to a much larger scale of variants through its access to natural genetic variants, or the WT alleleome (6), and with larger scales of data comes expectations of broader and deeper insights. In this study, we leveraged a very large set of variants from a comprehensive *E. coli* coding alleleome, comprising of variants from both ALE and WT coding alleles, to explore the solution space of all observed variants for tolerance to conditions of industrial importance.

Supplementing available ALE variants for genes of interest increased the total amount of unique variants to investigate by an order of magnitude, from hundreds to thousands of variants. Recent technologies enabled the reintroduction of this scale of variants in singleton (Onyx® Digital Genome Engineering platform). Selection experiments were executed on the pool of variants to understand each variant’s level of benefit according to their resulting enrichment. The selection experiments demonstrated that both ALE and WT contributed highly enriched variants, where the enrichment and benefits depended on the conditions, genes, and gene product regions. The results revealed that variants not originating from the same ALE experiment, or variants from outside the lab, can potentially confer substantially greater benefit.

Though there exist variants that have higher benefits than those of the original ICT ALE experiment (8) in singleton, ALE endpoint strains host multiple beneficial variants and likely reach higher levels of fitness. One could introduce multiple highly enriched variants for a condition into a strain, though interactions between variants, or epistasis, is also an important factor affecting the final fitness of the strain. Epistasis is likely already playing a role with the variants enriched in this study: there wasn’t complete agreement between the key variants of the ICT ALE studies and those highly enriched in this study (Figure 2C). There is evidence that at least some of the highly enriched variants of this study would be compatible together: the highly enriched variants were often found in the same areas of a gene as the original ICT ALE variants.

Variant trends on features of interest are a typical approach for interpreting the effects that variants are selected for (4). These trends often manifest as a clustering or overlapping of the beneficial characteristics of the variants. This study’s goal of maximizing variants investigated on genes of interest enabled easier interpretation of the possible beneficial changes variants were selected for. A broader study comparing the characteristics of WT and ALE variants found little overlap between their characteristics (6). This study also found little overlap between the AA positions of variants from different sources but did find valuable overlap with this and other characteristics. Variants from ALE and WT sources were often found in AA positions near each other, if not in the same position. Those variants of different sources found to overlap on the same position didn’t demonstrate generally higher enrichment than non-overlapping variants, though they were found in regions of a gene’s sequence that ultimately hosted highly enriched variants. The broader *E. coli* alleleome study found the coding alleleome is mostly conserved (6); the clustering or overlapping variants from different sources possibly represent areas of a gene that host an important variability in its functionality, which in turn can be leveraged as a genome design variable for applications of interest. Genes not hosting highly enriched truncations are hypothesized to be adjusting their functionality rather than completely removing it from the organism. The variants expected to accomplish these adjustments tend to demonstrate the highest enrichment with conservative rather than radical effects, suggesting even further constraints on the optimal solution space for genome design variables. Highly enriched variants from both variant sources also demonstrated an overlap of residue property changes across all case study genes. Property changes involving substantial residue side-chain size and flexibility changes were always highly-ranked and shared by variants from both sources in case studies, suggesting these property changes as primary mechanisms of functional changes for genes with this study’s conditions. These results indicate that the combination of variants from both ALE and WT variant sources better describes the solution space of genome design variables than only one of the sources.

This study describes the potential of using all observed variants to genes of interest towards identifying a beneficial set of variants for industrially relevant conditions. The initial set of observed variants possibly encapsulated a significant portion of the feasible functional solution space for genes and the phenotypes resulting from the selection experiments described each variant’s position on the fitness landscape. Understanding variant trends in relation to their targets and property changes offered valuable insights into how the molecular functions represented by the genes can be modulated for specific applications. It is anticipated that by analyzing increasing scales of variants from a variety of sources with data-driven strain design, we can continue to make progress toward more optimal application strain designs as well as fill in knowledge gaps with rational strain engineering methods.

## Methods

### Experimental Procedures

#### Bacterial Strains and Culture Conditions

Experiments were performed with the *E. coli* MG1655 strain and the cultures were grown in LB medium at 37 °C and 200 RPM when preparing the strain for genome editing with Onyx Digital Genome Engineering platform. The selection experiments were performed in M9 minimal media at 37 °C in growth profiler (Enzyscreen). Antibiotics are added to liquid cultures when appropriate.

Chemicals and antibiotics used include chloramphenicol (Merck), carbenicillin (Merck), 1,2-propanediol (Sigma-Aldrich), 2,3-butanediol (Acros organics), 1,6-diaminohexane (TCI), 1,4-diaminobutane dihydrochloride (TCI), glutaric acid (TCI), adipic acid (TCI), hexanoic acid (TCI), octanoic acid (Sigma-Aldrich), isobutyric acid (Sigma-Aldrich), p-coumaric acid (Sigma-Aldrich) and butanol (Sigma-Aldrich).

#### Library Design and Construction

Mutation libraries were constructed using high-throughput CRISPR-MAD7 based editing with Onyx Digital Genome Engineering Platform. Inscripta’s Designer Software was used to design gRNA and repair templates to create 7637 designs. The designer algorithm assigns design scores as high, mid, and low indicating the chances of successful edits. Redundant designs were used for the designs with mid and low scores to increase the probability of successful edits. Two plasmid system was used, one containing MAD7 enzyme and the second plasmid containing gRNA, repair template and unique barcode specific to each design. Editing cassettes were cloned in bulk into high-copy number plasmid backbone and plasmid populations were then electroporated into MAD-7 expressing cells using Inscripta’s reagents and in-built protocols in the onyx platform.

#### Sequencing of Library Samples

DNA (genomic DNA pool and plasmid DNA pool) was isolated from the cell library using Wizard SV DNA purification system (Promega) for determining genome engineering efficiencies. The library preparation protocol for edit identification assay and barcode diversity assay were performed according to Inscripta’s manual: https://inscripta.showpad.com/share/rWxQsFGmJznLKrdfWBHGH. The fraction of edited cells in the mutation libraries were determined using the edit-identification assay kit from Inscripta. Briefly, The genomic DNA pool was sequenced at 1000x coverage using Illumina’s MiSeq V2 2*150bp kit. The percentage of edited cells was determined to be 36% using Inscripta’s resolver software. Barcode diversity assay kit (Inscripta) was used to construct plasmid pool library for sequencing and sequenced using Illumina’s Miseq V2 2*150bp kit. Read1 fastQ data was uploaded to Inscripta’s Resolver Software to identify the relative abundance of each design in the library.

#### Selection Experiments

The library was initially enriched in half the final ALE concentration of 11 different industrial chemicals as the library did not grow when exposed directly to final ALE concentrations for 7 out of 11 chemicals. The final ALE concentration (9) of different chemicals used include 1,2-propanediol (83 g/L), 2,3-butanediol (79 g/L), hexamethyldiamine (38 g/L), putrescine (38 g/L), glutarate (47.5 g/L), adipate (50 g/L), butanol (16.2 g/L), isobutyrate (12.5 g/L), coumarate (20g/), octanoate (10 g/L) and hexanoate (7.5 g/L). Briefly, the library was inoculated at the starting OD of 0.08 in M9 medium containing 11 different chemicals, each in 3 replicates. The M9 medium without any chemical was used as control. In all cultures, antibiotics were added to maintain the plasmids. 96 w growth profiler plate and growth profiler from Enzyscreen was used for cultivation of all library samples and the growth was monitored in real-time. The library samples on coumaric acid and butanol did not grow even at the half the ALE concentration and thus eliminated from further selection. The enriched library from half the final ALE concentration was harvested in late-log or stationary phase by pelleting the cells. The cells were washed in M9 medium before inoculation to M9 medium with final ALE concentration of chemicals. The enriched cells at final ALE concentration of chemicals were harvested at stationary phase by pelleting the cells. Plasmid DNA pool was isolated from each replicate using automated plasmid extraction using MagJET plasmid extraction kit (ThermoFisher Scientific). Barcode diversity assay kit from Inscripta was used to construct libraries containing barcodes from plasmid pool for each sample for sequencing. The samples were sequenced using Illumina’s Nextseq midoutput 300 cycles kit. The fastQ data was uploaded to Inscripta’s resolver software to identify relative abundance of designs in each sample. The unique barcode for each design was used by the software to count the relative abundances of each sample.

#### Convergence Calculation

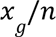

*x_g_* is the amount of replicate ALEs a gene *g* is mutated in for an ALE experiment, *n* is the total amount of replicate ALEs within an ALE experiment.

#### Enrichment Scores

Enrichment scores were calculated through Inscripta’s Differential Count Analysis (DCA) Tool (https://dca.sandbox.inscripta.com/), which uses the DESeq2 (19) package to analyze count data from different time points or under different conditions to test for differential representation. In gene editing experiments, typically the count data represents the counts or representation of a particular edit in an edit library.

#### Figure 2B Cutoffs

These were chosen according per condition to be between the median and upper quartile for enrichment scores across all genes of a condition. This enabled the analysis to better focus on genes hosting the most highly enriched variants.

#### Residue Properties

Residue properties were acquired from the software package *ssbio (20)*.

#### Application of Grantham Scores to Multi-AA Substitutions

For the broader analysis of Figure 1, Figure 2, and Figure S1, the highest Grantham scores across a multi-AA substitutions was kept to represent the whole variant. For case studies, Gratham scores were assigned to each AA within a multi-AA substitutions.

#### Gene Product Feature Annotations

Gene product features were acquired from EcoCyc (21), (22), Mutfunc (23), Interpro (24), and RCSB PDB (25).

#### Gene Structures and Visualizations

Protein structures were acquired from the AlphaFold Protein Structure Database (26,27). The protein structure of the visual abstract used NGL viewer (28).

#### Quantitative Plots

Plots were generated using Matplotlib (29) and Seaborn (30) Python software packages.

#### Software Scripts

The software scripts and data supporting the conclusions of this article are available at the following: https://github.com/biosustain/ale_v_wt_ict.git

#### Generative AI

ChatGPT, BARD, Claude.ai, and Grammarly were used to refine the narrative and grammar of this work, though were not used in the research process.

#### Availability of Data and Materials

The data sets supporting the conclusions of this article are available in the following open-access archive repository: https://github.com/biosustain/ale_v_wt_ict.git

#### Abbreviations

ALE: Adaptive Laboratory Evolution
WT: Wild-type
ICT: Industrial Chemical Tolerance
INDEL: Insertion or Deletion
AA: amino acid

## Author Contributions

Conceptualization and computational analysis: PVP. Conceptual support: BOP. Designed selection experiment, sequencing, and analysis, and troubleshot experimental data: VK. Executed selection experiment, and library preparation for sequencing: ZDJ. Technical support with WT variants: SMC. Writing: PVP, ZDJ, VK, BOP.

## Acknowledgments

The authors gratefully acknowledge Line Marcussen, Mariana Saavedra, and Kealan Peter Exley for their support with the experimental work.

