## supplemental information for "Using the *E. coli* Alleleome in Strain Design" (from another version of this preprint: version 1, 2023-09-17)

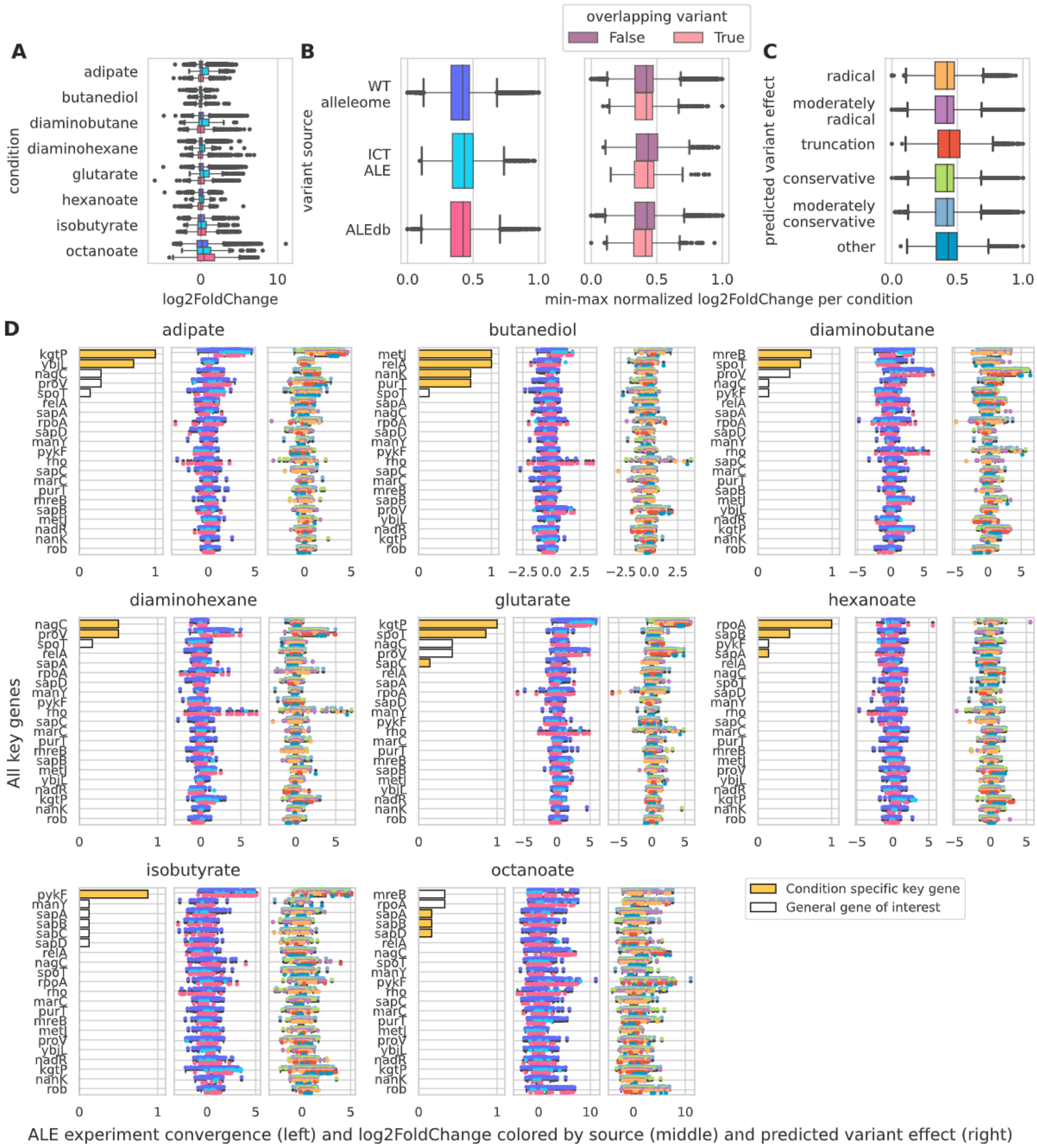


**Supplemental Figure 1** Distributions of variant enrichment on multiple levels. For non-normalized enrichment distributions, variants on the negative side of distributions enrich for the control of growth on glucose, while variants on the positive side of distributions enrich for growth on the treatment. **A)** Enrichment score distributions across conditions and variant sources. **B)** Normalized enrichment score distributions across conditions between variant sources and between amino acid substitutions found to or not to target amino acid positions seen in both variant sources. **C)** Normalized enrichment score distributions across conditions between predicted variant effects. **D)** Enrichment score distributions across conditions and key genes. ALE experiment convergence values indicate how many individual ALEs a gene were mutated in an ALE experiment. The color-to-source and predicted variant effect mappings are reused from subplots A, B, and C.

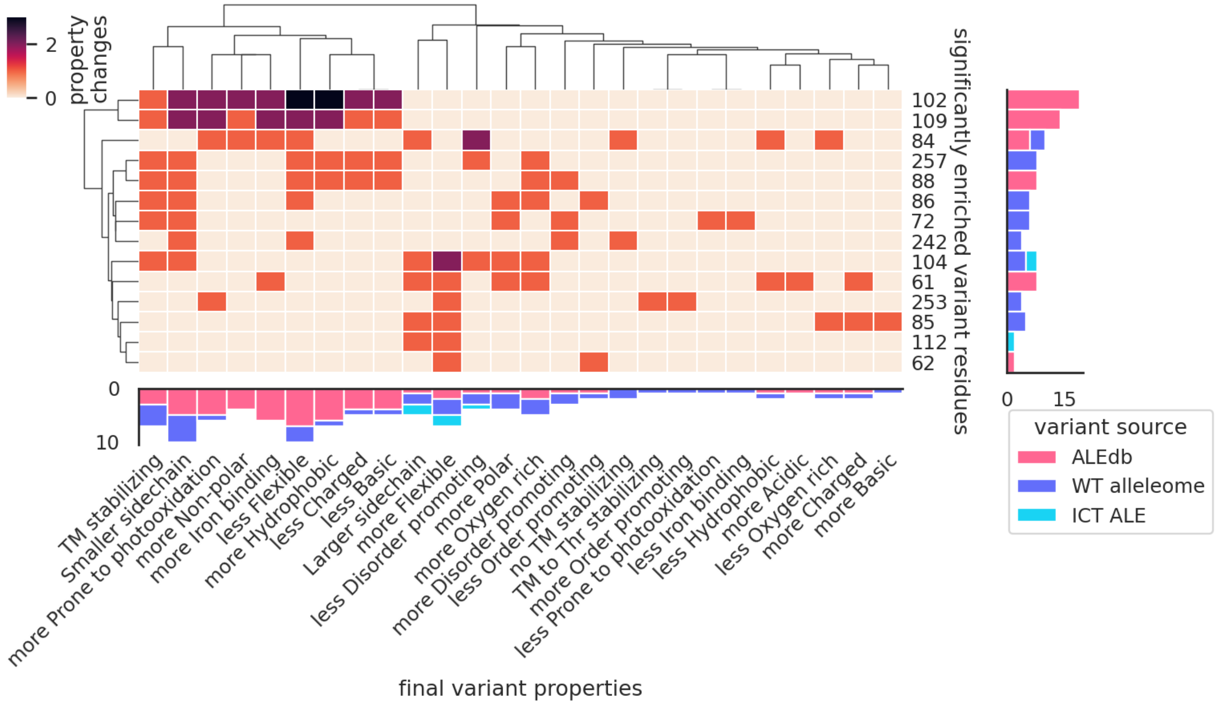


**Supplemental Figure 2** Clustering of property changes for highly enriched substitutions to Rho.

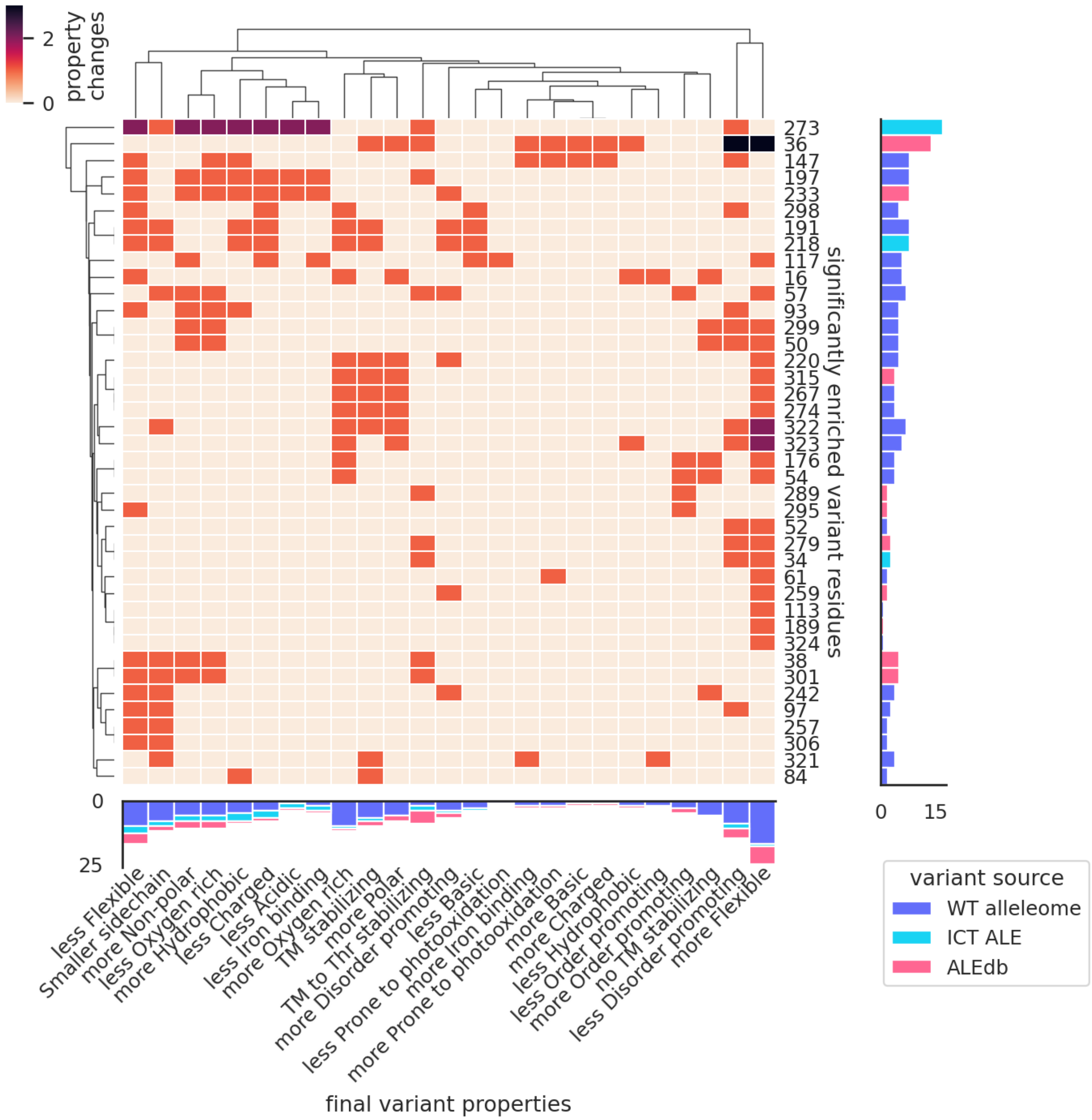


**Supplemental Figure 3** Clustering of property changes for highly enriched substitutions to RpoA.

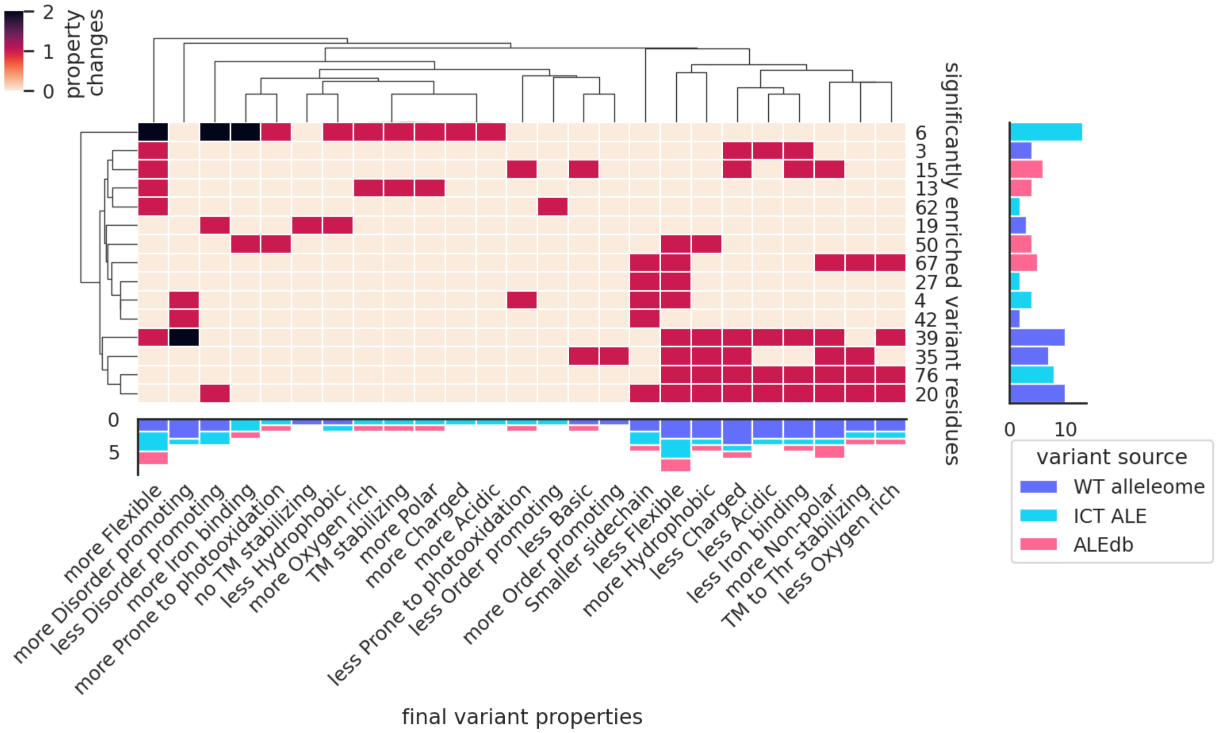


**Supplemental Figure 4** Clustering of property changes for highly enriched substitutions to MetJ.
